# Impact of impaired endogenous neurosteroidogenesis on outcomes following chronic alcohol exposure

**DOI:** 10.64898/2026.01.17.700089

**Authors:** Katrina Blandino, Yingchu He, Lifen Htet, Shadeh Okoudjou, Jonathan Lee, Maia Chinatti, Katie Ahn, Mike Lewis, Sarah Gray, Klaus Miczek, Jamie Maguire

## Abstract

Alcohol use disorder is a major public health concern worldwide and there is a high comorbidity with psychiatric disorders. The basolateral amygdala (BLA) has been implicated in both mood and alcohol use disorders; however, the mechanisms contributing to the shared pathophysiology remain unknown. Extensive evidence indicates that ethanol modulates GABAergic signaling in the BLA, including actions on neurosteroid-sensitive, extrasynaptic δ subunit-containing GABA_A_ receptors (GABA_A_Rs), which has been suggested to mediate many of the behavioral effects. In fact, several studies have suggested that 5α-reduced neurosteroids, such as allopregnanolone, may mediate some of the behavioral effects of alcohol. Here we demonstrate that chronic intermittent ethanol (CIE) exposure impairs endogenous neurosteroidogenesis via downregulation of key neurosteroidogenic enzymes, 5α-reductase type 1 and type 2. To examine the impact of impaired endogenous neurosteroidogenesis of the behavioral consequences of chronic alcohol exposure, including withdrawal-induced anxiety and increased alcohol consumption, we used CRISPR/Cas9 mediated knockdown of 5α-reductase in the BLA. Reduced expression of 5α-reductase in the BLA did not impact post-CIE alcohol intake or anxiety-like behaviors during withdrawal, perhaps because endogenous neurosteroidogenesis is already impaired following CIE. Therefore, we examined the impact of enhancing neurosteroid levels, treating mice post-CIE with SGE-516, a synthetic GABA_A_R positive allosteric modulator, which increased voluntary alcohol intake. These findings implicate endogenous neurosteroidogenesis in behavioral outcomes associated with withdrawal from chronic alcohol exposure. Further, this study suggests that targeting endogenous neurosteroidogenesis may be a novel and useful therapeutic target.

## 1. Introduction

Alcohol use disorder (AUD) is a prevalent public health concern with a high comorbidity with psychiatric disorders such as anxiety disorders and major depressive disorder. The COVID-19 pandemic has profoundly amplified the impact of alcohol on society, with multiple studies reporting increased drinking in the population due to a myriad of COVID-19 associated reasons such as stress, increased isolation, etc. (Pomazal et al. 2023, Stockwell et al 2021, Schmidt et al 2021, Gajdics et al 2023). Thus, there is a more urgent need for effective, evidence-based treatments to alleviate the impact of this disorder on society and individuals.

A significant barrier to recovery and abstinence is the negative symptoms associated with withdrawal. Alcohol withdrawal refers to the manifestation of negative symptoms that results from a period of abstinence, termed hyperkatifeia (Wise and Koob 2014). Current treatments of AUD combine behavioral strategies with medications (Tareen et al 2024); however, current treatments do not directly address the underlying pathophysiology of disease and the negative symptoms associated with withdrawal. GABA_A_ modulators have been considered as treatment options, but there are concerns due to the fact that some of these treatments are associated with severe side effects and have their own abuse potential (Gatta et al 2022). Thus, there is a need for a better understanding of the neurobiological mechanisms contributing to the comorbidity between stress, psychiatric illnesses, and AUD with the goal of developing more effective, safer treatments, which specifically address negative affective states associated with withdrawal from chronic alcohol use.

A GABA_A_ receptor modulator that shows therapeutic potential is allopregnanolone, a neurosteroid synthesized either de novo from cholesterol or from the metabolism of progesterone, a process involving the rate-limiting enzyme, 5α-reductase (Paul et al. 2020). Allopregnanolone is a positive allosteric modulator (PAM) of the GABA_A_ receptor, acting preferentially at extra synaptic δ subunit-containing GABA_A_ receptors (GABA_A_Rs) with limited side effects and abuse potential. Brexanolone and zuranolone, which both have GABA PAM activity, have received FDA approval for the treatment of postpartum depression. Clinical trials with brexanolone, a proprietary formulation of allopregnanolone, demonstrated that in addition to prolonged reduction in depression symptoms, anxiety symptoms show extended improvement (Meltzer-Brody et al. 2018, Epperson et al 2023). Preclinical studies demonstrated that allopregnanolone has therapeutic benefit for many of the brain alterations associated with alcohol withdrawal such as inflammation, decreased GABA_A_ signaling and excess CRF (Morrow et al. 2020, Morrow et al. 2024). The blunting of these effects may be risk factors for developing AUD (Morrow et al. 2006). These studies beg the question whether GABA PAMs may be useful for combatting the negative affective states associated with alcohol withdrawal. In preclinical models, expression of 5α-reductase (type 1 and 2) RNA and protein increase in the prefrontal cortex of rats with a history of chronic and acute ethanol exposure (Sánchez et al 2014); whereas 5α-reductase expression has been shown to be decreased in the basolateral amygdala (BLA) following chronic stress exposure (Walton et al. 2023). Adding to the complexity, allopregnanolone and ethanol have been shown to influence one another (Sanna et al. 2004, Ford et al. 2008). There is evidence that neurosteroids, including allopregnanolone, also have an impact on ethanol signaling and sensitivity (VanDoren et al. 2000). Further, studies also demonstrated that administration of allopregnanolone directly to the nucleus accumbens and ventral tegmental area reduce ethanol self-administration (Ornelas et al. 2023). These rodent studies demonstrate a bidirectional relationship whereby ethanol exposure influences allopregnanolone and 5α-reductase expression, and the activity of these molecules also impacts ethanol related behaviors and signaling in the brain (Khisti et al. 2002). Thus, neurosteroids sit at the intersection between alcohol and emotional processing.

Clinical studies also support a relationship between allopregnanolone and alcohol exposure. Circulating allopregnanolone levels are elevated in response to acute intoxication and 5α-reductase expression patterns are associated with behaviors like alcohol craving (Torres and Ortega 2003, Lenz et al. 2012). Allopregnanolone may also be associated with the reward pathways involved in alcohol exposure. However, other studies show lowered allopregnanolone in plasma associated with alcohol withdrawal (Romeo et al. 2000). Thus, more studies are needed to understand the mechanisms through which allopregnanolone interacts with stress and alcohol to understand the role of endogenous neurosteroid synthesis in contributing to AUD and the therapeutic potential of drugs that leverage this biology.

The BLA is a brain region where alcohol consumption and stress intersect. Stress is strongly associated with BLA activity and BLA hyper-excitability is associated with anxiety disorders in humans and animal models. Studies have found that BLA hyper-excitability is associated with AUD severity (Sharp 2017). Studies have also observed dysregulated glutamatergic connections between the BLA and prefrontal cortex after exposure to chronic ethanol (Crofton et al. 2022). Chronic stress has been shown to impact allopregnanolone synthesis in the BLA and a reduction of 5α-reductase in the BLA was sufficient to induce behavioral deficits reminiscent of chronic stress (Walton et al. 2023). Importantly, over-expression of 5α-reductase was able to prevent the behavioral deficits associated with chronic stress exposure, suggesting that targeting endogenous neurosteroidogenesis may have therapeutic potential (Walton et al. 2023). As the BLA sits at the intersection between alcohol and stress, we sought to investigate how endogenous neurosteroidogenesis in this area may influence outcomes following chronic alcohol exposure.

It is still unknown what specific role allopregnanolone synthesis in the BLA plays in mediating anxiety, particularly in the context of alcohol withdrawal. The goal of this study is to investigate how neurosteroid synthesis in the BLA affects anxiety and drinking during withdrawal. Utilizing a vapor model of chronic ethanol exposure, the study investigates the impact of chronic ethanol on neurosteroid synthesis in BLA and how impairing synthesis affects anxiety behaviors in withdrawal. Further, this study explores how treatment with a GABA PAM (SGE-516) after exposure affects drinking behaviors in withdrawal. This study lends useful insight into the role that allopregnanolone metabolism, treatment with GABA PAMs, and GABA_A_R signaling play in anxiety in withdrawal and explores its therapeutic potential.

## 2. Materials and Methods

### 2.1 Animal Handling and Husbandry

Male and female C57Bl/6J animals (Strain #000664) and constitutive ROSA::Cas9-EGFP animals (male and female) were purchased from the Jackson Laboratory (Strain #026179) at 8-12 weeks of age. All mice were housed in a 12-hour light/dark cycle with ad libitum access to food and water. Surgical procedures were completed when animals were 11-13 weeks old. All animals were group housed except during the two-bottle choice drinking paradigm. Three days after the last Chronic Intermitted Ethanol (CIE) exposure in SGE-516 experiments, C57Bl/6J animals were single housed and assigned to receive either regular chow or SGE-516 chow (Teklad Rodent Chow #7012, 450mg SGE-516/kg). The chow assignment was counterbalanced across the four vapor chambers. All procedures were completed with approval from the Tufts University IACUC.

### 2.2 Surgical Procedures

Constitutive Cas9 animals were anesthetized with a cocktail containing 100mg/kg ketamine and 10mg/kg xyalzine. Sgsrd5α1 and sgsrd5α2 viruses (Walton et al. 2023) containing an mCherry tag and were mixed in a 1:1 ratio and 250nL of virus was stereotaxically injected into the BLA using the following coordinates: AP-1.5, ML±3.3, and DV-5.1 at a rate of 100nL/min using a 33-gauge Hamilton syringe (Hamilton 80100, 7803-05 syringe) and stereotaxic pump (WPI Instruments Smart Touch Micro 2T and UMP3 arm). The control virus AAV-hysn-mCherry was administered at the same volume and rate in control animals. Animals were given 0.5mg/kg buprenorphine slow release for analgesia and monitored for three days post-op. Animals were allowed to recover for one week for viral expression before experimentation. Knockdown of 5α-reductase type 1 and/or 2 (5αR) was verified by qRT-PCR. Misses, confirmed by no mCherry expression and no reduction in the expression of 5α-reductase 1 and 2, which were not significantly different than mCherry viral controls, were included as controls.

### 2.3 Chronic Intermittent Ethanol (CIE)

Vapor chambers were acquired from La Jolla Alcohol Research Institute (Model #TM112). Animals were randomly assigned to CIE or air conditions. Constitutive Cas9 animals were assigned to air and vapor groups such that they were counterbalanced based on the average of the last two weeks of baseline intermittent access drinking. Before starting chronic intermittent ethanol (CIE) exposure, all animals were injected with 1mmol/kg pyrazole (Sigma Aldrich #P56607-5G) and animals assigned to the CIE group were given a 1.6g/kg loading dose of ethanol (Rodberg et al. 2017, denHartog et al. 2020, Becker and Lopez 2004). Blood ethanol concentration (BEC) was maintained between 180-200dL/mL. Animals were exposed to ethanol vapor for 16 hours per day, Monday-Thursday from 5pm-9am for two consecutive weeks. In between vapor sessions animals were given 0.3mL subcutaneous saline, diet recover gels (Clear H_2_O, 72-07-5022), clear hydrogels (Clear H_2_O, 70-01-1062) and were maintained on heating pads. Intoxication was measured via the rotor rod after removal from the vapor chambers.

### 2.4 Behavioral Assessment of Intoxication

#### a. Rotor Rod

Upon removal from vapor chambers, animals were placed on a rotor rod (Maze Engineers) to assess coordination after vapor exposure (Blum et al. 1971, Krahe et al. 2017). Animals were placed on six lanes and tested at a maximum of 35rpm at an acceleration of 6rpm^2^. After thirty minutes, if animals had not fallen, they were removed from the rotor rod, and their latency was recorded as 1800 seconds. For each animal, latency to fall (seconds), distance traveled (cm) and speed at fall (rpm) were recorded.

### 2.5 Behavioral Assessments of Withdrawal

#### a. Social Interaction Test

The impact of ethanol on social avoidance was assessed in a three chambered apparatus (López-Cruz et al. 2016, Raymond et al. 2019, Simon et al. 2023). Animals were given one hour to habituate to the behavior room prior to the start of experiments. Chambers were wiped with 70% ethanol or chlorine dioxide prior to the start. Animals were placed in a three-chamber apparatus for a three-minute habituation period with two empty pencil cups on either side of the apparatus (three chambers 30 x 30 cm each). For the social interaction trial, a sex-matched younger novel animal was placed under a pencil cup on the right or left and the test animal was placed back in the chamber for three minutes. Animals were tested for social behavior four times throughout the protocol, (before Intermittent Access (IA) in the cas9 animals, before CIE, after one week of CIE, and after the final CIE) and the side of a novel animal alternated each time and alternated between animals. Behavior was recorded and scored using Ethovision computer software. Behavior was measured as the ratio of the time spent in the social chamber (the side with the novel animal) to the time in the non-social chamber (the side with the empty cup) in the social interaction trials.

#### b. Open Field Test

Avoidance behavior was tested following CIE using an open field task (Neira et al. 2022, Crabbe Jr. et al. 1982, Kliethermes 2005, Acevedo et al. 2014). Twenty-four hours after their last exposure to CIE vapor, animals were individually placed into the center of the open field apparatus and behavior was automatically scored using Motor Monitor II software throughout the ten-minute test. Animals were counterbalanced by treatment and sex in order tested. Behavior was measured as the ratio of time in the center relative to the time in the periphery of the open field apparatus.

#### c. Light/Dark Box Test

Exploration in the light/dark box is an established measure of avoidance behavior in rodents (Kliethermes 2005, Acevedo et al. 2014). Forty-eight hours after their last exposure to ethanol vapor, animals were placed in the dark half of the apparatus to start, and the distance, time, and entries into the light chamber were automatically measured using Motor Monitor II software throughout the ten-minute test. The testing order of animals was counterbalanced by treatment and sex. Behavior was measured as the ratio of time in the light side to the dark side of the light/dark box.

#### d. Nesting Behavioral Assessment

Nesting was used as measure of motivated behavior (Greenberg et al 2016). Mice were introduced to clean cages with a fresh nestlet measured for starting weight. Twenty-four hours later, the remaining nestlet was weighed. Behavior was measured as the percent of the nest that was unshredded after twenty-four hours.

### 2.6 Intermittent Access (IA) Two Bottle Choice Test

Constitutive Cas9 and C57Bl/6J animals were single housed in small Ancare mouse cages (AN75 Mouse bottoms) and provided with bottles filled with either 20% ethanol or water. Bottles were alternated to prevent a place preference and control drip cage was used to measure leakage or evaporation. Bottles were provided for a twenty-four-hour period Monday-Tuesday, Wednesday-Thursday and Friday-Saturday and were replaced with two new water bottles on off days. Drinking was measured as g/kg/24hrs, as well as preference for ethanol (ethanol consumed divided by the total liquid consumed). The average loss of fluid in drip cages was subtracted from either the water or ethanol totals for each animal.

### 2.7 5_α_-reductase Expression

#### a. Brain Dissection

Animals were anesthetized with isoflurane and rapidly decapitated. Brain tissue was rapidly fixed in 4% paraformaldehyde, cryoprotected in sucrose, and stored in 4°C for target validation using mCherry immunofluorescence imaging. For RT-qPCR and western blotting of 5α-reductase expression in the BLA, tissue was snap frozen in liquid nitrogen and stored in -80°C until experimentation. Frozen brains were sectioned on a cryostat (Leica) and a biopsy needle was used as a tissue punch to remove the basolateral amygdala. Fixed non-frozen brain tissue was rinsed in 1x PBS and sectioned on a vibratome (Leica VT1000S) at 50µm thick sections. They were stored in 1x PBS at 4°C covered in foil. Slices were mounted on Superfrost Plus Microscope Slides (Fisherbrand #1255015) with VECTCASHIELD HardSet Antifade Mounting Medium with DAPI (Vector Laboratories Inc. H-1500-10). Slides were cover slipped (Epredia Coverslips No.1 thickness #102460). Autofluorescence of the mCherry tag attached to the virus was imaged using a Keyence confocal microscope (Keyence #BZ-X710).

#### b. qRT-PCR Measurements

RNA extraction:

RNA was extracted by adding 25µL TRIzol (Invitrogen 15596018) per sample and then samples were homogenized using a sonicator. After sitting at room temperature for three minutes, 15µL of chloroform was added and samples were vortexed and centrifuged for 15 minutes at 4°C 12 x g. The clear chloroform layer was separated into prepared tubes of 2µL glycogen (Thermo Scientific RO551); samples were spun a second time to maximize recovered sample. 12.5µL of cold isopropanol was added to each sample and they were vortexed and allowed to sit at room temperature for ten minutes. They were then spun at 12 x g for 45 minutes at 4°C. The supernatant was then removed, and the pellet was resuspended in 500µL of cold 80% ethanol. The samples were dislodged and briefly spun at 7.5xG for 5 min at 4°C. The ethanol was removed, and the pellets were allowed to dry. The samples were resuspended in 20µL of cold 1mMol sodium citrate (RNASecure, Invitrogen Thermo Scientific AM7000) and were placed on a heat block at 60°C for 15 min and then briefly vortexed and spun down. The concentration (ng/µL) and purity (260:280) ratio were measured using a Nanodrop (ThermoFisher). Samples were stored in -80°C.

qRT-PCR: 100ng of template RNA was used for RT-qPCR experiments with the following primer sets:

Srd5α1:5’-GAG-ATA-TTC-AGC-TGA-GAC-CC-3’(Forward)

5’-TTA-GTA-TGT-GGG-CAG-CTT-GG-3’ (Reverse)

Srd5α2: 5’-ATT-TGT-GTG-GCA-GAG-AGA-GG-3’ (Forward)

5’-TTG-ATT-GAC-TGC-CTG-GAT-GG-3’ (Reverse)

beta actin: 5’-GGC-TGT-ATT-CCC-CTC-CG-3’ (Forward)

5’-CCA-GTT-GGT-AAC-AAT-GCC-ATG-T-3’ (Reverse)

Transcript levels for 5α1 (Srd5α1) and 2 (Srd5α2) were quantified using a SuperScript III SYBR Green Dye Platinum One-Step qRT-PCR kit (Thermo Invitrogen # 11732088) with an Mx3000 (Stratgene) and StepOnePlus (Thermofisher) cyclers using the following cycle parameters: 1 cycle of 50 °C 3 min, 1 cycle 95 °C 5 min. 40 cycles of 25 sec of 95°C and 30 sec 60°C , 1 cycle 1 min 40°C and 1 cycle of 1 min 95°C , 30 sec 55°C and 30 sec 95 °C. Change in CT values relative to beta actin was measured (Darnieder et al. 2019, Walton et al. 2023, Evans-Strong et al. 2024) and analyzed for each animal with naïve animals (novel animals used in social interaction) used as controls for handling stress.

#### c. Western Blotting

Protein was extracted in 200µL of a lysis buffer containing 1x RIPA buffer (Boston BioProducts, Inc. BP-115X), 20% PMSF dissolved in 100% ethanol (10µL per 1mL of RIPA buffer) and 1 tablet of EDTA-free proteinase inhibitor (Roche Diagnostics #11836170001). Samples were homogenized on ice with a sonicator and left on ice for 1hr. Concentration of the protein was measured using a DC assay (BioRad #500-0116). 100µg of protein was separated by gel electrophoresis and transferred to a PVDF membranes at 4°C at 100V for one hour. Membranes were then blocked in 10% milk in 1x TBST overnight. Blots were incubated with primary antibodies for 5α1 and 2 (Abcam goat anti-mouse srd5α1 # ab110123 and 2 #ab27469) in 5% milk in 1x TBST at a concentration of 1:1000 for two hours. Secondary antibodies (Jackson Immuno Research Donkey anti-goat HRP #705-035-003) were mixed in 5% milk at a concentration of 1:2000 and incubated for two hours. Membranes were then developed using Tanon High-Sig ECL Western Blotting Substrate (Cat#180-5001) and visualized on a Bio-Rad Universal Hood III (# 610-01). Blots were re-probed for beta tubulin as loading control using a primary antibody against beta tubulin (Sigma Monoclonal anti-beta tubulin produced in mouse) at a 1:1000 concentration and secondaries (GE Healthcare ECL Sheep anti-Mouse IgG Horseradish peroxidase linked whole antibody) at a 1:2000 concentration. Protein intensity was measured using Image J and normalized to b-tubulin expression.

### 2.8 Tissue Measurements of Allopregnanolone and BEC

#### a. Submandibular Blood Draws

Blood was collected via the submandibular vein to measure allopregnanolone in plasma at the following timepoints: before the first IA drinking period, before CIE, after the first week of CIE, after the second week of CIE and trunk blood was collected at time of sacrifice. In experiments testing SGE-516 vs control chow, samples were taken before post-CIE voluntary drinking (1 week after first exposure to SGE-516/control chow). Blood was also drawn on days four and eight for BEC concentration after vapor exposure. Animals selected were counterbalanced between cohorts for sex and vapor chamber. Blood was collected in CAT serum lined (Greiner Bio One 450470) or heparin lined (Greiner Bio One 450477) collection tubes. They were spun down at 4°C for 30min at 1.8x g speed and the plasma layer was removed and stored at -80°C.

#### b. BEC Measurements

Samples for BEC measurements were drawn from one control (Air) and one vapor exposed mouse (CIE), randomly selected, once per week for BEC measurements. Ethanol percent in blood was measured using EnzyChrome colorimetric assay (EnzyChrome, ECET-100) according to manufacturer’s instructions. Measurements were read using a plate reader at 595nm and concentrations were determined against a standard curve.

#### c. Plasma and BLA Brain Punch Allopregnanolone Measurements

Allopregnanolone was measured in the plasma using the Arbor Assays DetectX Allopregnanolone ELISA kit (Product K061-H5) according to manufacturer’s instructions. Samples were then stored in 1x assay buffer at -20°C. For brain allopregnanolone measurements, lysates were collected from brain samples using the methods described above for western blotting and allopregnanolone was extracted using the liquid steroid extraction method from the protocol for Arbor Assays DetectX Allopregnanolone ELISA kit (Product K061-H5). Samples were stored in 1x Assay Buffer in -20°C until assayed. Allopregnanolone measurements were read at 450nm and calculated based on a standard curve.

### 2.9 Data Analysis

Data were analyzed via GraphPad Prism and Microsoft Excel software using a Students unpaired t-test. Paired t-tests or 2-Way ANOVAs were used where appropriate. Outliers were identified by the ROUT test (Q=1) in Prism and excluded from the data analysis. 5αR knockdown animals that were identified as misses had a difference from the control virus average greater than zero in both Srd5α1 and 2 and were included with control virus animals for analysis. Animals with a single miss were considered knockdowns and data was included with the double knockdown animals. 5α-reductase knockdown animals that were lost to acute withdrawal mid-CIE were not included in baseline analyses. Data are reported as the mean±SEM.

## 3. Results

### 3.1 Chronic Ethanol Exposure Induces Behavioral Deficits Associated with Withdrawal

Two weeks of CIE vapor were sufficient to induce behavioral deficits associated with intoxication and withdrawal in C57Bl/6J mice (Figure 1). Immediately following CIE exposure, intoxication levels were measured using a rotor rod test, demonstrating decreased latencies to fall, consistent with intoxication, in CIE mice compared to Air controls (Air: 453.8±75.33sec., CIE: 81.87±12.47sec., n= 10 Air, 8-12 CIE, Mixed Effects Model, Time x Tx History, Effect of Tx History F(1,20)=21.92, p=0.0001, Supplemental Figure 1A). Mice subjected to CIE exhibit an increase in avoidance behaviors in the open field test measured as a ratio of time spent in the center versus the periphery compared to control mice exposed only to air (Air 0.1317±0.01892, CIE: 0.06132±0.00797, n= 18 Air, 17 CIE, unpaired t-test, t(33)=3.355, p=0.002, Figure 1B). Similarly, mice subjected to CIE exhibit increased avoidance behaviors in the light/dark box, spending a decreased amount of time in the dark chamber compared to the light (Air: 1.042,± 0.1271; CIE: 0.5294 ± 0.08122, n = 18 Air, 16 CIE, unpaired t-test, t(32)=3.302, p=0.0024, Figure 1B). There was also increased nesting behavior observed in animals exposed to CIE, measured by the percent nest unshredded (Air: 44.5±10.36%, CIE: 20.08±5.036%, n=12 Air, 12 CIE, unpaired t-test, t(22)=2.120, p=0.0455, Figure 1B). Animals also show a trend towards increased drinking in the IA two-bottle choice paradigm following CIE (Air: 12.24±1.092 Avg. g/kg/24h s., CIE: 15.5±1.310 Avg. g/kg/24hrs., n= 11 Air, 15 CIE, unpaired t-test, t(24)=1.817, p=0.0818, Figure 1C). These data confirm the behavioral correlates of withdrawal following chronic alcohol exposure in mice.

**Figure 1.**
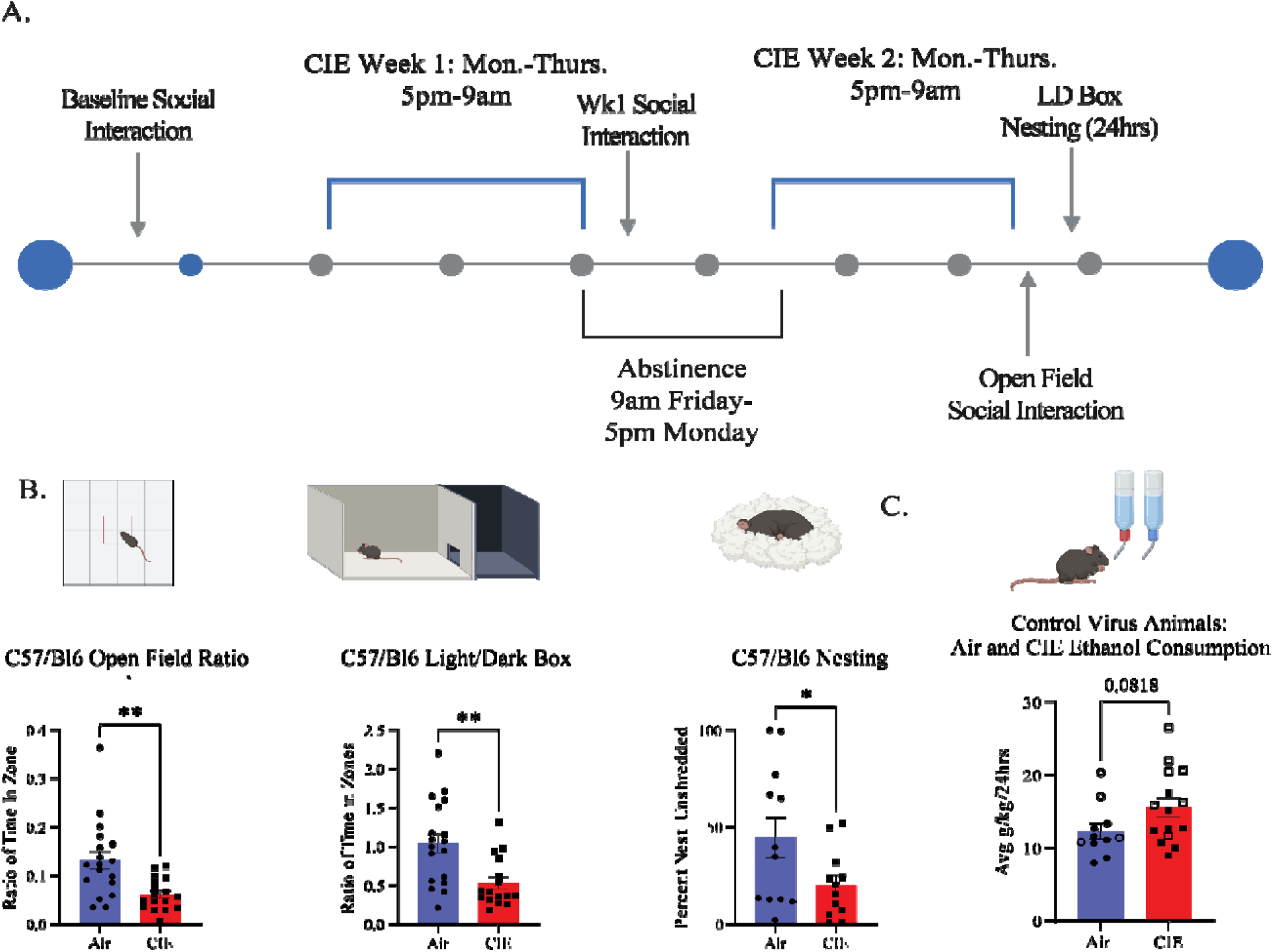
CIE exposed C57/Bl6 male mice show higher avoidance behaviors than air control mice after two weeks of exposure. A.Timeline of CIE vapor exposure over two weeks with avoidance behavior testing. B. Ratio of time in center of open field to the periphery 24hrs. post-CIE (left, unpaired t-tests, p=0.0020, n=18 Air, 18 CIE), time in the light to the dark in light/dark box 48hrs. post-CIE (center, unpaired t-test, p=0.0024, n=18 Air, 16 CIE) and percent unshredded nest 48-72 hrs. post-CIE (right, unpaired t-test, p=0.0455, n=12 Air, 12 CIE). C. Average ethanol consumption of mCherry injected cas9 mice exposed to air or CIE (unpaired t-test, p=0.0818, n=7 male Air, 4 female Air,8 male CIE, 7 female CIE) Males are in closed symbols and females in open symbols. Data are presented as the mean±SEM.

### 3.2 Chronic Ethanol Exposure Impairs Endogenous Neurosteroidogenesis

To explore the role of endogenous neurosteroidogenesis in the behavioral deficits observed following CIE, we measured allopregnanolone levels in whole brain tissue and plasma (Figure 2A). Allopregnanolone levels were not significantly different in the plasma following CIE compared to Air control (Air: 4201±971.7pg/mL; CIE: 10794± 3391 pg/mL; n= 8 Air, 9 CIE, unpaired t-test, t(15)=1.771, p=0.0969, Figure 2B), and there was no difference in the ratio of total brain allopregnanolone to plasma levels (Air: 9.519±3.905, CIE: 10.49±3.488, unpaired t-test, p=0.8614, Figure 2C). To examine potential changes in the capacity for endogenous neurosteroidogenesis in the brain following CIE, we examined 5α-reductase expression in the BLA. One week following CIE exposure, there was a significant reduction in the protein expression of 5α-reductase type 1 compared to air control (Air 1.005 ± 0.06545; CIE: 0.4014± 0.02986 n =3 Air, 3 CIE, unpaired t-test, t(4)=8.391, p=0.0011) with no observable change in transcript expression (Air: 0.6544±0.05794; CIE: 0.5306 ± 0.09527; n =30 Air, 10 CIE, unpaired t-test, t(38)=1.081, p=0.2866, Figure 2D-E). These data suggest changes in the capacity for endogenous neurosteroid synthesis following CIE.

**Figure 2.**
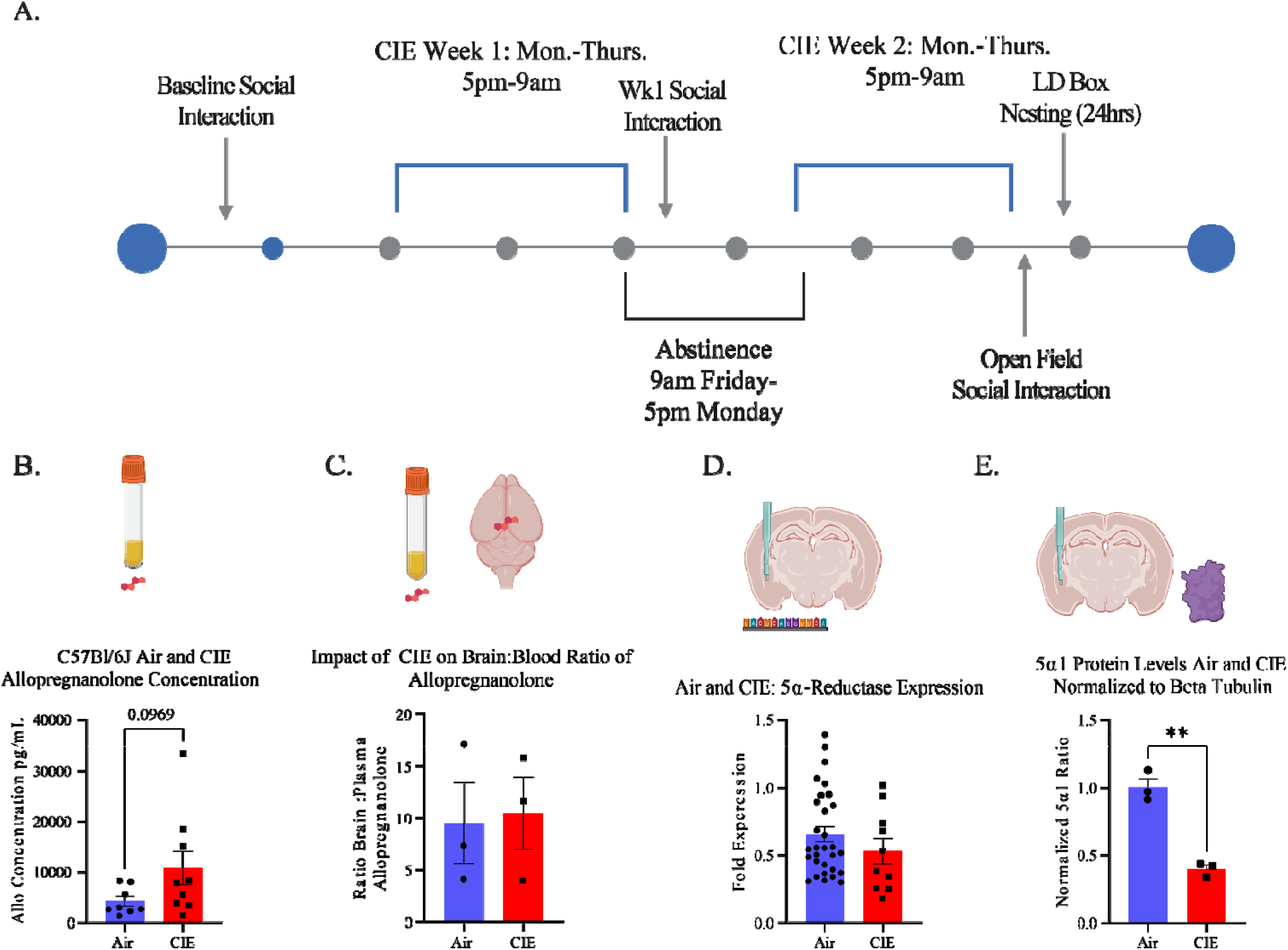
CIE exposed C57/Bl6 mice show a decrease in 5_α_1 expression in the BLA after 1 week of exposure to CIE. A. Timeline of CIE vapor exposure over two weeks. B. Plasma allopregnanolone in C57Bl/6J mice after CIE exposure (n=8 Air, 9 CIE, unpaired t-test, t(15)=1.771, p=0.0969).C. Ratio of whole brain allopregnanolone of animals to plasma allopregnanolone in air and CIE animals (unpaired t-test, p=0.8614, n=3 Air, 3 CIE).D. RNA expression of 5_α_1 and 2 of male and female animals in the BLA after CIE (unpaired t-test of 5_α_1 and 2 expression values, p=0.2866, n=30 Air, 10 CIE). E. 5_α_-reductase type 1 expression relative to beta tubulin in male animals exposed to 1 week CIE (unpaired t-test, p<0.0001, n=3 Air, 3 CIE). Data are presented as the mean±SEM.

### 3.3 Impaired Endogenous Neurosteroidogenesis Does Not Impact Baseline Alcohol Intake

To examine the impact of a reduced capacity for allopregnanolone synthesis on alcohol intake and withdrawal behaviors, we used a CRISPR::Cas9-EGFP mouse line to selectively knockdown 5_α_-reductase in the BLA using guide RNAs targeting 5_α_-reductase type 1 and 2, sgsrd5_α_1 and sgsrd5_α_2 respectively. This approach was sufficient to knockdown the expression of 5_α_-reductase as measured by difference from control virus mean expression of 5_α_R1 (control: 3.957±3.43, 5αR1 knockdown: -5.978±0.3537, n=12 control virus, 9 5αR1 knockdown, One Sample T test, theoretical mean 0.0, control: t(11)=1.154, p=0.2731, 5αR1 knockdown t(8)=16.9, p<0.0001) and 5αR2 (control: 0.3896 ±0.7576, 5αR2 knockdown:-1.654±0.05132, n=11 control virus, 9 5αR2 knockdown, one sample t-test, theoretical mean 0.0, control t(11)=0.5143 p=0.6182, 5αR knockdown t(8)=34.88, p<0.0001). These findings are similar to previous reports using this method (Walton et al. 2023).

Interestingly, knockdown of 5α-reductase expression in the BLA did not alter baseline drinking levels in the IA two-bottle choice paradigm in males (Control: 9.449±0.6668 g/kg/24hrs., 5αR knockdown: 9.358±0.6962 g/kg/24hrs.; n = 20 Control, 18 5αR knockdown, unpaired T test, t(36)=0.9256, p=0.9256) or in females (Control: 16.06±1.444 Avg. g/kg/24hrs., 5αR knockdown: 13.97±1.619 Avg g/kg/24hrs., n=12 Control, 10 5αR knockdown, unpaired t-test, t(20)=1.012, p=0.3236) (Figure 3D). Similar to numerous previous reports, we do observe an increase in voluntary alcohol intake in females compared to males in Control (Males: 9.449±0.6668g/kg/24hrs.; Females: 16.06±1.444 g/kg/24hrs.; n = 20 males, 12 females, unpaired t-test, t(30)=4.703, p<0.0001) and 5αR knockdown animals (Males 9.358±0.6962 Avg. g/kg/24hrs., Females 13.97±1.619 Avg. g/kg/24hrs., n= 18 males, 10 females, unpaired T test, t(26)=3.263, p=0.0031, Supplemental Figure 3A).

**Figure 3.**
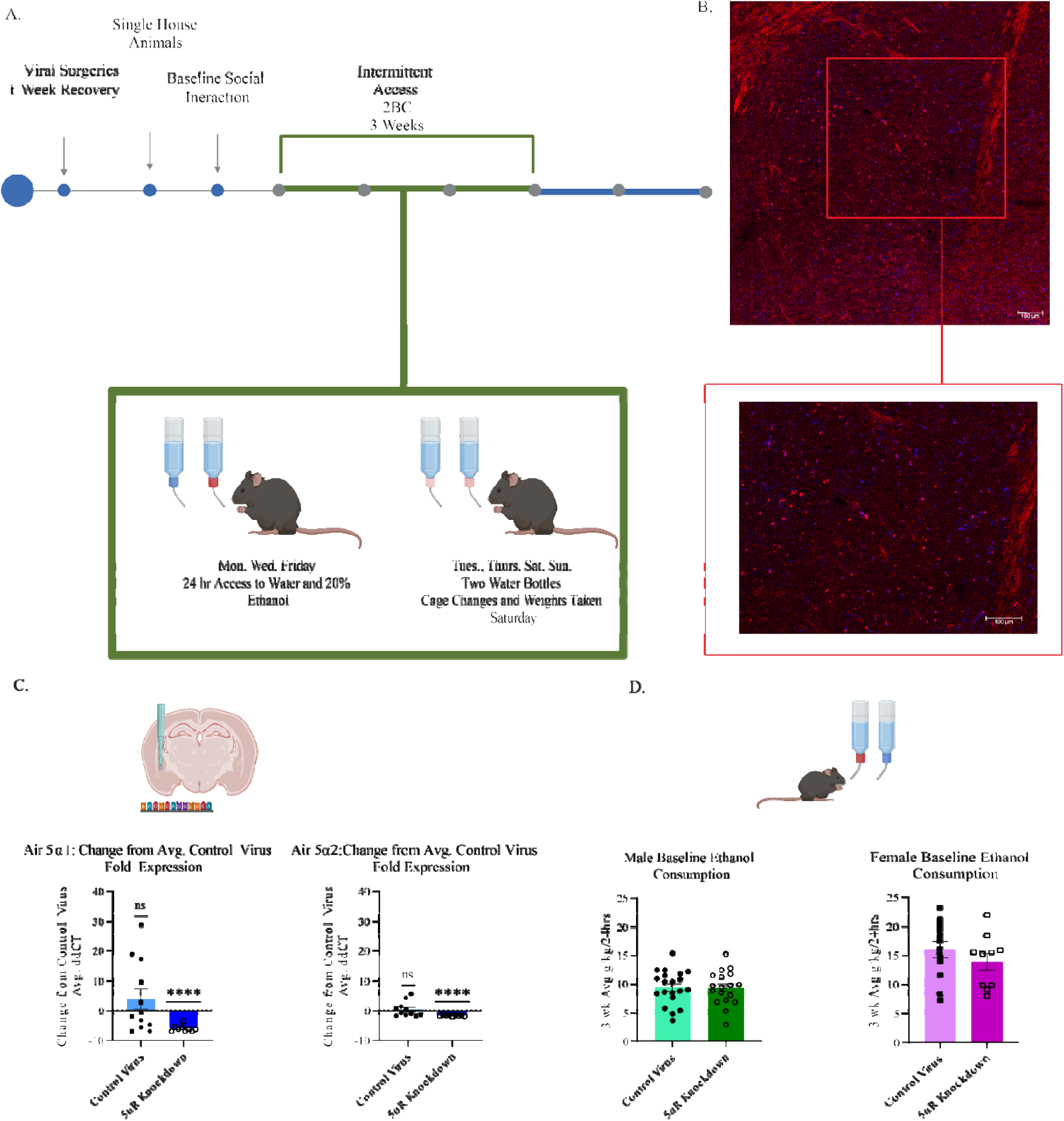
5_α_1- and 5_α_2- reductase expression is reduced in 5_α_-reductase knockdown animals and there is no impact of the 5_α_-reductase knockdown on baseline drinking. A. Experimental timeline showing surgery and baseline intermittent access drinking before CIE. B. Representative image (40x) of mCherry tag on sgsrd5_α_1/2 virus injection site in basolateral amygdala with higher magnification showing cellular expression. C. Change from control virus average ddCT fold change in Air exposed control and 5α-reductase (5αR) knockdown males (circles) and females (squares) for 5α1 (left, n=12 control virus, 9 5αR knockdown, one sample t-test, theoretical mean 0.0, p=0.2731, 5αR knockdown, p<0.0001) and 5α2 (right, n=11 control virus, n=9 5αR knockdown, one sample t-test, theoretical mean 0.0, control virus p=0.6182, 5αR knockdown, p<0.0001). D. Baseline (pre-CIE) ethanol consumption in males (left, n=20 Control, n=18 5αR knockdown, unpaired t-test, p=0.9256) and females (right, n=12 Control, n=10 5αR knockdown, unpaired t-test, p=0.3236). ns= not significant, \*\*\*\**p* < 0.0001. Data are presented as the mean±SEM.

### 3.4 Impaired Endogenous Neurosteroidogenesis Does Not Impact Behavioral Deficits Associated with Withdrawal

The impact of impaired endogenous neurosteroidogenesis on withdrawal-related behaviors was examined in the 5αR knockdown mice following CIE. To measure intoxication, we measured latency to fall from the rotor rod immediately after time in the vapor chambers and, consistent with the C57BL/6J mice, we see decreased latency to fall in CIE control virus males relative to air (Air: 271.3±20.01 sec., CIE: 168.7 ± 17.16 sec., n=10 Air, 8 CIE, Mixed Effects Model, Tx History x Time, Effect of Tx History F(1,16)=18.35, p=0.0006) and control virus females relative to air (Air: 345.3±33.37sec., CIE: 210.3±15.78sec., n=4 Air, 8 CIE, 2-Way ANOVA Tx History x Time, Effect of Tx History F(1,10)=5.187, p=0.0460, Supplemental Figure 1B). Interestingly, while we see an effect of CIE to reduce latency in 5αR knockdown CIE mice relative to air in males (Air: 287.5±38.38sec., CIE: 168.6±20.33sec., n=7 Air, 12 CIE, Mixed Effects Model, Tx History x Time, Effect on Tx History F(1,17)=13.69, p=0.0018), there were no differences in female 5αR knockdown animals between air and CIE (Air: 268.2±17.38sec., CIE: 248.4.4±19.21 sec., n=4 Air, 6 CIE, 2Way ANOVA Tx History x Time, Effect of Tx History F(1,8)=0.2700, p=0.6174, Supplemental Figure 1C).

There were no differences in avoidance behaviors in the open field test measured as the ratio in the center versus periphery in 5αR knockdown males (Control: 0.03119± 0.005964; 5αR knockdown: 0.03783± 0.004563) (n=8 Control, 12 5αR Knockdown, unpaired t-test, t(18)=0.8974, p=0.3814) or females (Control: 0.04846± 0.008049; 5αR: 0.04236 ± 0.009746) (n=8 Control, 6 5αR knockdown, unpaired t-test, t(12)=0.4861, p=0.6357, Figure 4B). Similarly, there were no differences in avoidance behaviors in the light/dark box measured as the ratio of time in the light compartment compared to the dark in males (Control: 0.2261 ± 0.03770; 5αR knockdown: 0.2118 ± 0.01732), (n=8 Control, n=10 5αR knockdown, unpaired t-tests, t(16)= 0.3704, p=0.7159) (Figure 4B) or females (Control: 0.2838± 0.04085; 5aR knockdown: 0.4622 ± 0.1210; n = 8 Control, 6 5αR knockdown, unpaired t-test, t(12)= 1.568, p=0.1428) (Figure 4B).

**Figure 4.**
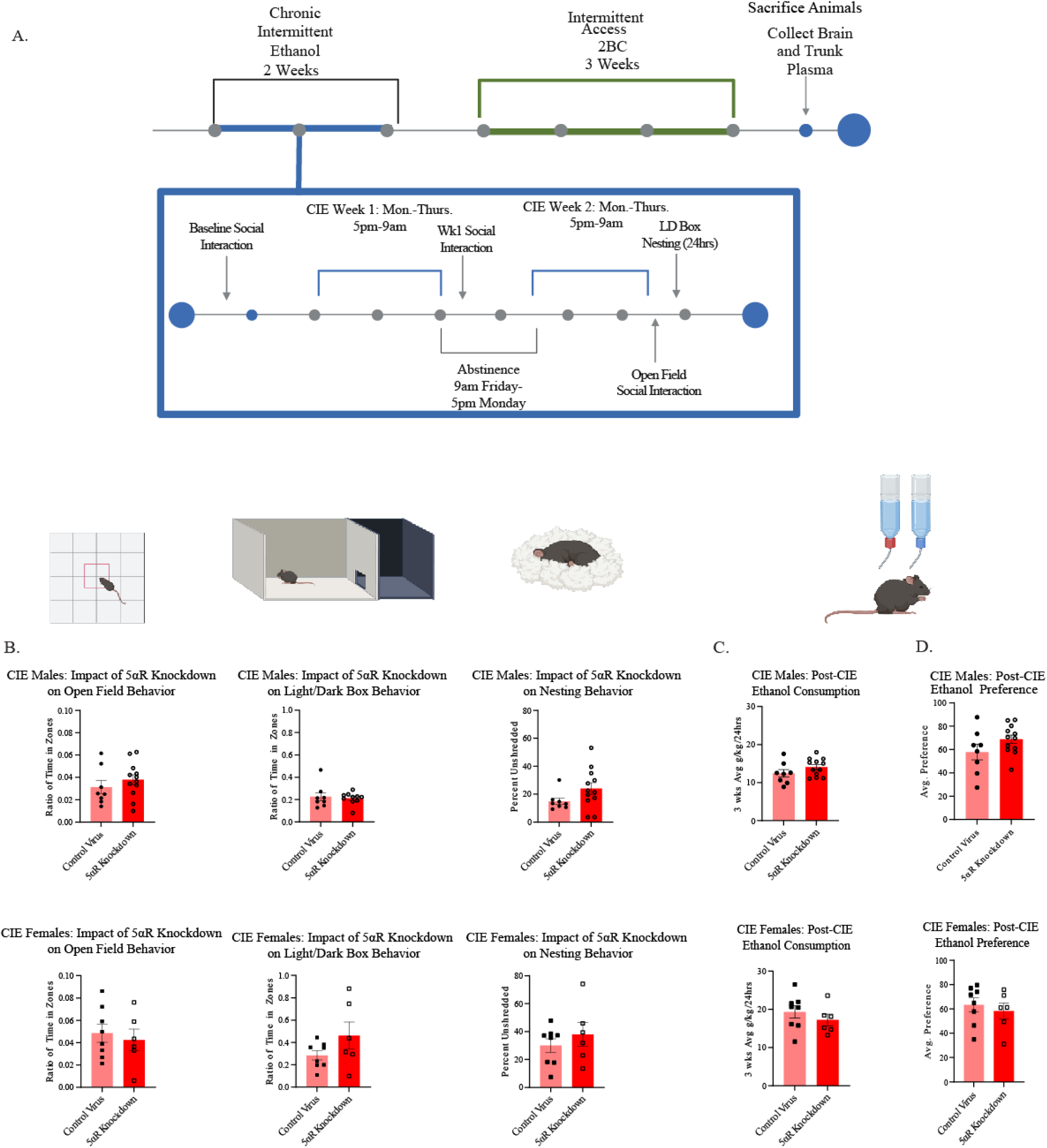
There is no impact of 5_α_-reductase knockdown in the BLA on post-CIE avoidance behaviors, or on ethanol consumption. A. Experimental timeline showing CIE exposure timeline and post-CIE drinking. B. Avoidance behaviors of CIE exposed males (top) and females (bottom) in the open field 24hrs. post exposure (upper left: males n=8 Control, n=12 5αR Knockdown, unpaired t-test, p=0.3814, lower left: females n=8 Control, 6 5αR knockdown, unpaired t-test, p=0.6357), the light/dark box 48hrs post-CIE (upper center: males n=8 Control, 10 5αR knockdown, unpaired t-tests, p=0.7159, upper center: females n=8 Control, 6 5αR knockdown, unpaired t-test, p=0.1428) and nesting 48-72hrs. post exposure (upper right: males n=8 Control, n=12 5αR knockdown, unpaired t-test, p=0.1067, upper right: females n=8 Control, n=6 5αR knockdown, unpaired t-test, p=0.4086). C. Post-CIE drinking in CIE exposed male (upper, n=8 control virus, n= 12 5αR knockdown animals, unpaired t-test, p=0.1733) and female animals (lower, n=8 control virus, n=6 5αR knockdown, unpaired t-test, p=0.3781) . D. Ethanol preference post-CIE in CIE exposed male (upper, n=8 control virus, n=12 5αR knockdown, unpaired t-test, p=0.1414) and female animals (lower, n=8 control, n=6 5αR knockdown, unpaired t-test, p=0.5698). Data are presented as the mean±SEM.

There were no changes in nesting behavior in 5αR relative to controls in males (Control: 14.84±2.325%, 5αR knockdown: 24.06±4.123%, n=8 Control, 12 5αR knockdown, unpaired t-test, t(18)=1.698, p=0.1067) and females (Control: 29.96±4.855%, 5αR knockdown: 37.92± 8.626%, n=8 Control, 6 5αR knockdown, unpaired t-test, t(12)=0.8563, p=0.4086) in the nestlet shredding test (Figure 4B). We measured social behavior as the change in percentage time spent in the social chamber each week of CIE relative to the pre-IA baseline social behavior (Supplemental Figure 2A). Social behavior does not change from baseline in control or 5αR knockdown male mice in weeks 1 (Week 1 control virus: - 2.026±5.626 difference in percent time, 5αR knockdown:-4.480±7.872 difference in percent time, n=5 control virus, 9 5αR knockdown, one sample t-tests, Theoretical Mean 0.0, control virus t(4)=0.3602, p=0.7369, 5αR knockdown t(8)=0.5692, p=0.5849) or week 2 of CIE (Week 2 control virus: 0.8575±8.818 difference in percent time, 5αR knockdown: 8.208±7.404 difference in percent time, n=5 control virus, 9 5αR knockdown, one sample t-tests, Theoretical Mean 0.0, control virus t(4)=0.09725, p=0.9272, 5αR knockdown t(8)=1.109, p=0.2998, Supplemental Figure 2B). Similarly, female control virus or 5αR knockdown mice show no change from baseline in CIE week 1 (Week 1 control virus: 1.772±9.739 difference in percent time, 5αR knockdown: 6.524±6.983% difference in percent time, n=8 control virus, 6 5αR knockdown, One Sample t-tests, Theoretical Mean 0.0, control virus t(7)=0.1819, p=0.8608, 5αR knockdown t(5)=0.9344, p=0.3930) or CIE week 2 (Week 2 control virus: 0.03609±8.724 difference in percent time, 5αR knockdown: -2.385±13.08 difference in percent time, n=8 control virus, 6 5αR knockdown, one sample t-tests, Theoretical Mean 0.0, control virus t(7)=0.004137 p=0.9968, 5αR knockdown t(5)=0.1823, p=0.8625, Supplemental Figure 2C).

Impaired endogenous neurosteroid synthesis also does not impact ethanol consumption following CIE in males (control virus: 12.44±0.9926 Avg. g/kg/24hrs., 5αR knockdown: 14.07±0.6676 Avg. g/kg/24hrs., n=8 control virus, 12 5αR knockdown animals, unpaired t-test, t(18)=1.418, p=0.1733) or females (control virus: 19.33±1.635 Avg. g/kg/24hrs., 5αR knockdown: 17.22±1.521 Avg. g/kg/24hrs., n=8 control virus, 65αR knockdown, unpaired t-test, t(12)=0.9152, p=0.3781) (Figure 4C). Ethanol consumption was not altered either in air exposed 5αR knockdown animals (Males: 11.82 ±1.217 Avg. g/kg/24hrs., Females 15.13±2.246 Avg. g/kg/24hrs.) relative to control virus animals (Males:11.57±0.7908 Avg. g/kg/24hrs., Females: 14.84±2.321 Avg. g/kg/24hrs.) (Males n=9 control virus, 7 5αR knockdown, unpaired t-test, t(14)=0.1793, p=0.8603, Females n=4 control virus, 4 5αR knockdown, unpaired t-test, t(6)=0.08892, p=0.9320, Supplemental Figure 3B).

Preference for ethanol is not altered either in CIE exposed males (control: 57.92±6.728%, 5αR knockdown 68.71±3.611%, n=8 control virus, n=12 5αR knockdown, unpaired t-test, t(18)=1.538, p=0.1414) or females (control: 63.33±5.606%, 5αR knockdown: 58.28±6.5966.827%, n=8 control, n=6 5αR knockdown, unpaired t-test, t(12)=0.5844, p=0.5698, Figure 4D). Preference was not altered either by impaired neurosteroidogenesis in air 5αR knockdown males (57.87±5.547%) or females (53.58±5.688%) relative to air exposed control virus animals (Males: 56.25±4.978%, Females: 50.81±7.099%, Males n=9 control, n=7 5αR knockdown, unpaired t-test, t(14)=002170, p=0.8313, Females: n=4 control, 4 5αR knockdown, unpaired t-test, t(6)=0.3051, p=0.7706, Figure 3C). While there are no changes in ethanol consumption or preference due to impaired neurosteroid synthesis, animals escalate drinking as expected after CIE exposure in the control animals (3.152±0.7191 g/kg/24hrs., n=16, One Sample T test, Theoretical Mean 0.0, t(15)=4.383, p=0.0005) and CIE exposed 5αR knockdown animals (3.690±0.6604 g/kg/24hrs., n=18, one sample t-test, Theoretical Mean 0.0, t(17)=5.587, p<0.0001). This escalation was not seen in air exposed control virus (0.7849±1.202 g/kg/24hrs., n=13, one sample t-test, Theoretical Mean 0.0, t(12)=0.6531, p=0.5260); yet interestingly there is escalation of drinking in air exposed 5αR knockdown animals (3.310±1.284 g/kg/24hrs., n=11, one sample t-test, Theoretical Mean 0.0, t(10)=2.579, p=0.0275, Supplemental Figure 3D). This is likely due to the stress of the paradigm. Thus, while the impact of CIE on chronic withdrawal is present, the knockdown of 5α-reductase does not further induce behavioral deficits following CIE, perhaps because endogenous neurosteroidogenesis is already impaired.

### 3.5 Behavior in Acute Withdrawal Correlates with Allopregnanolone in Knockdown Females Following CIE

To measure the role of allopregnanolone in mediating avoidance behaviors in withdrawal, we correlated plasma allopregnanolone concentrations to performance on behavioral tests at week 2. There were no observed correlations between plasma at the second week of CIE and avoidance behavior in the open field in males (Control: n=7, r(5)=-0.6429 p=0.1389, 5αR knockdown: n=11, r(9)=-0.1455, p=0.6731, Spearman Correlation, Figure 5A). A correlation is observed in 5αR knockdown females but not control females (Control: n=8, r(6)=-0.07143, p=0.8820, 5αR knockdown n=6, r(4)=-0.9429, p=0.0167) (Figure 5B). Thus, there may be compensatory changes in peripheral allopregnanolone synthesis in the face of impaired neurosteroid synthesis in the brain associated with behavioral changes following CIE.

**Figure 5.**
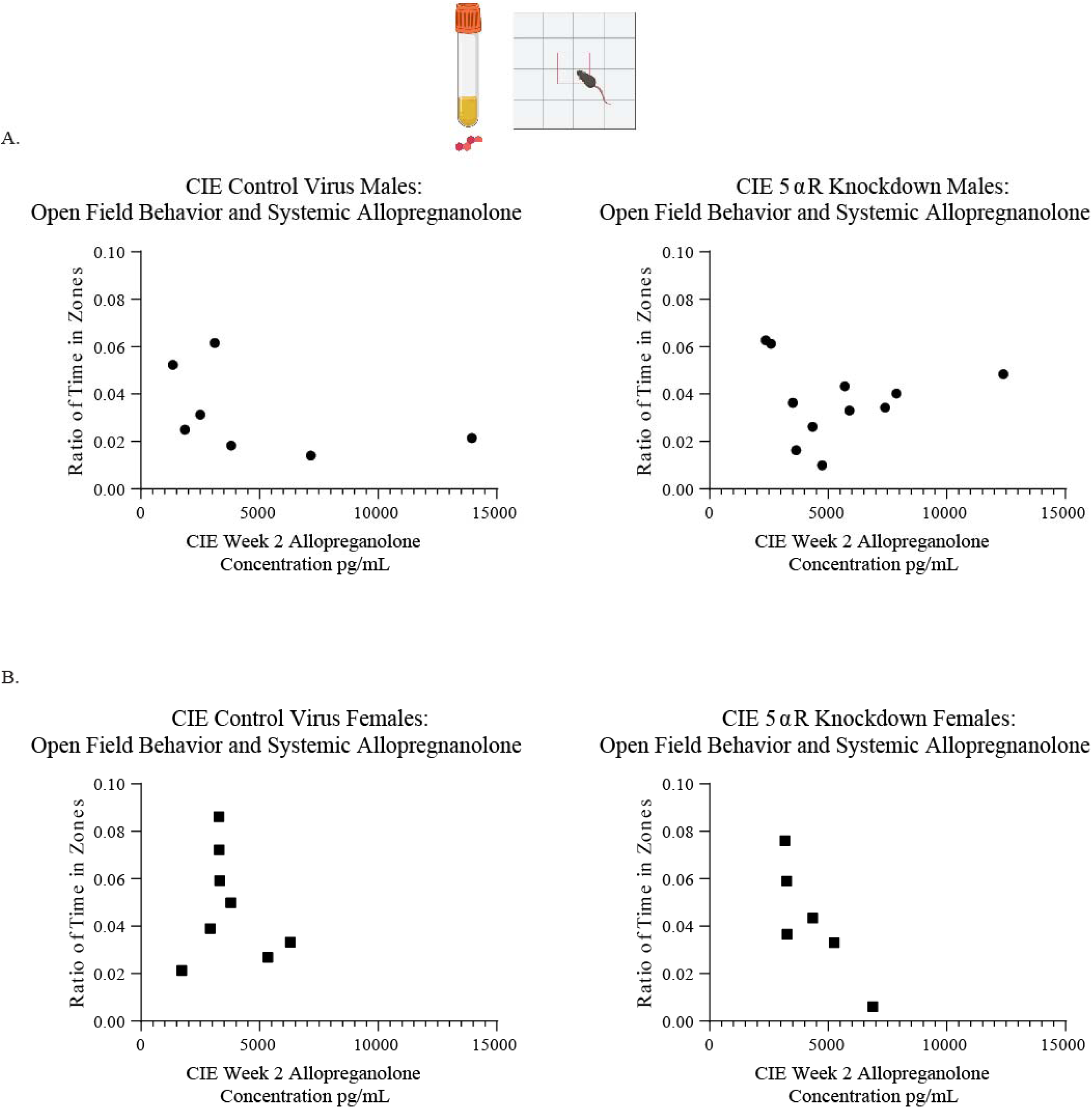
There is a negative correlation between plasma allopregnanolone and open field avoidance behavior in CIE exposed 5_α_R knockdown females. A. Correlation between avoidance behavior in the open field and plasma allopregnanolone concentration in CIE exposed male control virus (left, n=7, Spearman Correlation, p=0.1389) and 5αR knockdown animals (right, n=11, Spearman Correlation, p=0.6731). B. Correlation between avoidance behavior in the open field and plasma allopregnanolone concentration in CIE exposed female control virus (left, n=8, Spearman Correlation, p=0.8820) and 5αR knockdown animals (n=6, Spearman Correlation, p=0.0167).

### 3.6 Treatment With a GABA PAM Increases Alcohol Intake Following CIE

To directly test the impact of neurosteroid-based treatments on increased alcohol consumption associated with withdrawal following CIE, we treated mice with SGE-516, a GABA PAM tool compound developed by SAGE Therapeutics, and evaluated the impact on post-CIE voluntary alcohol intake (Figure 6A). Surprisingly, we observe an increase in voluntary alcohol consumption in the IA two-bottle choice paradigm following CIE in males (14.31±0.6865 Avg g/kg/24hrs.) and females (17.49 ± 0.5138 Avg. g/kg/24hrs.) compared to standard chow-treated male (8.5± 2.093 Avg. g/kg/24hrs.) and female mice (14.17 ±1.034 Avg. g/kg/24hrs.) (Males: n=4 regular chow, 5 SGE-516 chow, unpaired t-test, t(7)=2.911, p=0.0226, Females: n=8 regular chow, 7 SGE-516 chow, unpaired t-test, t(13)= 2.749, p=0.0166, Figure 6B). We verified that SGE-516 chow did not alter BLA concentrations of allopregnanolone compared to regular chow in males (regular chow: 1501±242.0pg/mg, SGE-516 chow: 1844±391.8pg/mL/mg, n= 4 regular chow, 5 SGE-516 chow, unpaired t-test, t(7)=0.6965, p=0.5086) or females (regular chow: 1371±157.1pg/mL/mg, SGE-516 chow: 1213±137.9pg/mL/mg, n=8 regular chow, 8 SGE-516 chow, unpaired t-test, t(14)=0.7585, p=0.4608, Supplemental Figure 4A ).

**Figure 6.**
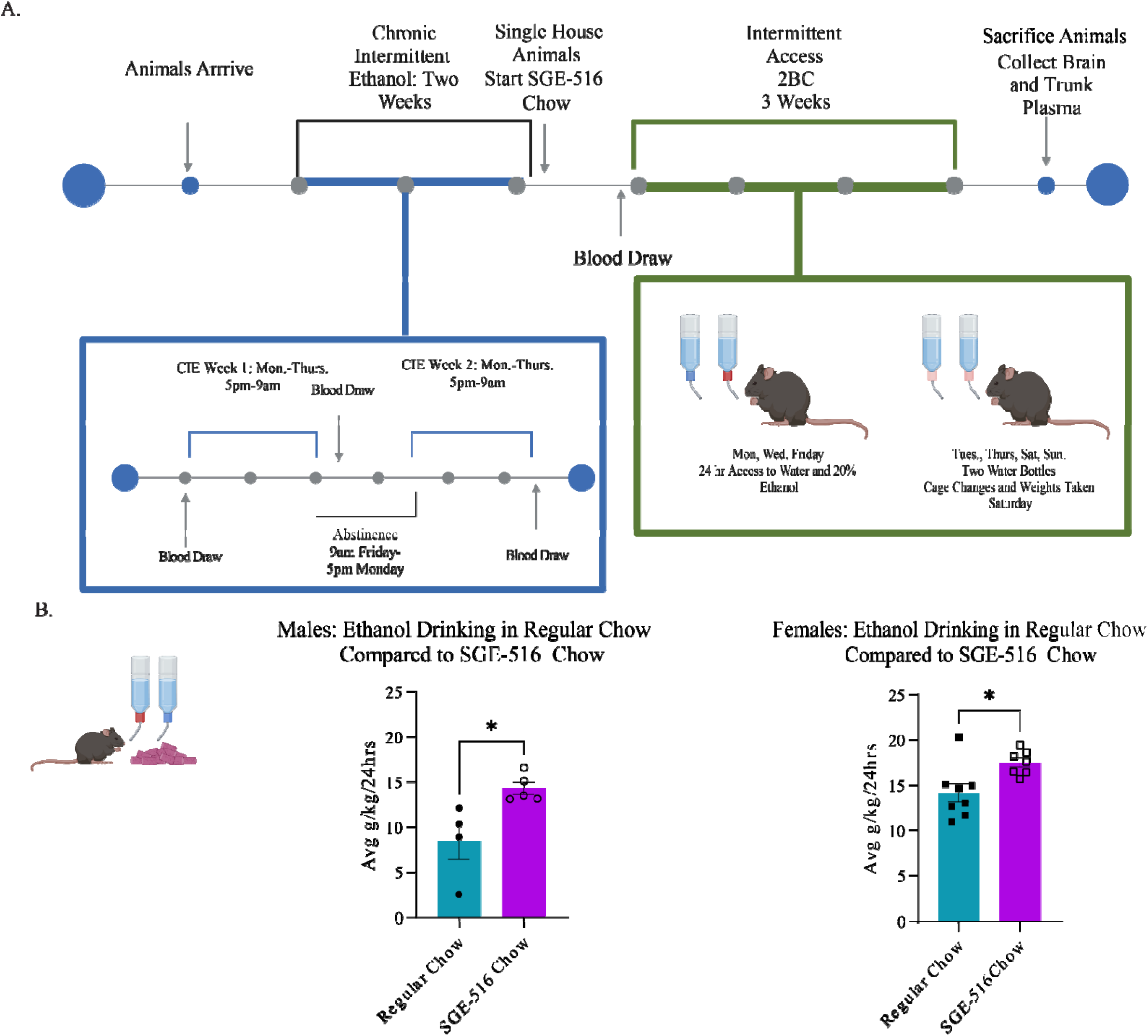
Animals fed SGE-516 chow drink more than regular chow animals. A. Experimental timeline showing CIE exposure, SGE-516 chow feeding and intermittent access drinking. B. Average of three weeks of drinking in males (top) and females (bottom) fed regular chow and SGE-516 chow (unpaired t-, males: n=4 regular chow, 5 SGE-516 chow, p=0.0226, females: n=8 regular chow, 7 SGE-516 chow, p=0.0166). Data are presented as the mean±SEM. \**p* <0.05.

Similar to our previous reports (Walton et al. 2023), we demonstrate that SGE-516 treatment is capable of altering 5α-reductase expression, increasing 5α-reductase transcript expression in male mice fed SGE-516 chow (2.785 ± 0.2133 fold 5αR expression) relative to regular chow males (1.611±0.1131 fold 5αR expression, n=4 regular chow, 5 SGE-516 chow, unpaired t-, t(7)=4.489, p=0.0028, Supplemental Figure 4), suggesting that GABA PAMs may impact endogenous neurosteroid synthesis. There was no impact of SGE-516 on 5α-reductase transcript levels in the BLA in female mice (regular chow: 2.633±0.2926, SGE-516: 2.356±0.3359, n=7 regular chow, 8 SGE-516, Unpaired T test, t(13)=0.6132, p=0.5503, Supplemental Figure 4B). These data are consistent with the influence of allopregnanolone on alcohol intake.

## 4. Discussion

The objective of this study was to determine the role of endogenous neurosteroid synthesis, particularly allopregnanolone synthesis via 5α-reductase, in the BLA on behavioral outcomes following exposure to chronic ethanol. We observe the expected increased avoidance behavior in wildtype C57Bl/6J mice exposed to CIE vapor relative to air controls (Figure 1B), consistent with the negative affective states accompanying withdrawal from alcohol. We demonstrate that wild type animals show a reduction in 5α-reductase type 1 expression in the BLA following CIE, which we proposed may contribute to the behavioral deficits associated with alcohol withdrawal. However, CRISPR/Cas9-mediated knockdown of 5α-reductase in the BLA did not worsen the behavioral outcomes following CIE (Figure 4B), potentially because 5α-reductase expression is already reduced by CIE leading to a floor effect. Similarly, knockdown of 5α-reductase did not alter alcohol consumption or preference following CIE (Figure 4C-D). These data are consistent with previous reports demonstrating that 5α-reductase expression in the BLA sets a baseline affective tone (Walton, et al. 2023) and decreased expression following CIE may contribute to the behavioral deficits associated with withdrawal from alcohol. Overall, these data are also consistent with previous reports suggesting that neurosteroids mediate the pharmacological effects of alcohol such as the sedative, anti-convulsant, and anxiolytic effects of ethanol as well as the impact of ethanol on spatial learning (Matthews et al. 2002, Khisti et al. 2002). These data demonstrate that more work is needed to understand the dynamics between levels of 5α-reductase expression in the context of ethanol exposure and the impact on ethanol withdrawal associated behaviors

In light of these findings, we investigated the therapeutic potential of exogenous neurosteroid-based treatments on the increased alcohol intake following CIE exposure. Treatment with SGE-516, a GABA PAM, increased alcohol intake in both males and females. The increased voluntary alcohol consumption following CIE in SGE-516 treated animals was associated with an increase in 5α-reductase expression in the BLA. in males These findings contradict our expected results, but once again demonstrate that neurosteroids have the potential to influence alcohol intake. Previous studies show conflicting results when examining the impact of exogenous allopregnanolone on ethanol consumption. While some studies have demonstrated that administration of acute allopregnanolone before drinking sessions elevated female mice drinking while eliciting a dose-dependent effect in males (Finn et al. 2010), others show that administration of allopregnanolone to the nucleus accumbens and ventral tegmental area reduce drinking in rats (Ornelas et al. 2023). Studies also suggest an impact of dose on the effects of allopregnanolone on ethanol consumption; a middle range dose increased self-administration for ethanol while a high dose of allopregnanolone caused reduced responding (Janak et al. 1998). In fact, a recent study demonstrated that a high dose of allopregnanolone decreased binge-like alcohol consumption, but treatment with the GABA PAM, zuranolone, only transiently (within hours) reduced alcohol intake (Ursich, 2026). Thus, the chronic nature of exposure to SGE-516 chow during drinking in our study may maintain a dose of GABA PAMs that facilitates alcohol intake. It is possible that impaired endogenous neurosteroidogenesis in the 5αR knockdown mice leads to elevated alcohol intake over time by reducing the rewarding effects of ethanol, leading to increased alcohol intake; whereas exogenous neurosteroid treatment (SGE-516) may enhance the effects of alcohol (or make them more tolerable) leading to increased alcohol intake. Further studies are required to fully understand the mechanisms through which neurosteroids influence alcohol intake associated with withdrawal.

Alcohol toxicity and neuroinflammation are well-documented consequences of chronic breakdown of alcohol in the brain. Long term exposure to ethanol results in increased cytokine activation, chronically stressing the immune system of the brain (Pascual et al. 2018). Rodents exposed to ethanol after brain injury showed cognitive impairment in addition to enhanced reactivity in astrocytes and microglia (Teng et al. 2015). Neuroinflammation has also been associated with anxiety-like behaviors within two weeks; increased microglia activation was observed in the amygdala of rats that displayed anxiety-like behaviors while recovering from injection of neuraminidase (León-Rodríguez et al. 2022). Recent work also highlights how ethanol exposure suppresses microglia in the BLA of female mice and male mice that have been exposed to stress and ethanol. Stress elevated microglia activation in female mice, and in chronic withdrawal, male mice show this elevation in activated microglia (Soares et al. 2025). Female mice also showed altered ethanol place preference after injury, an effect that was mediated by sex steroids (Oliverio et al. 2022). This highlights the role of time scale in how neuroinflammation may be impacting behavior as well as sex-specific steroids.

Neuroinflammation has already been identified as a mechanism through which neurosteroids influence alcohol (Morrow et al. 2024). Allopregnanolone has been shown to inhibit toll-like receptors (Balen et al. 2021) and given the pro-inflammatory effects of alcohol, this is a likely mechanism mediating the effects of neurosteroids on alcohol intake. This study focused on how the disruption in endogenous neurosteroid synthesis during withdrawal may alter the effects of neurosteroids on alcohol intake compared to non-dependent animals. Thus, neurosteroid-based treatments may still offer therapeutic benefit for AUD outside of acute withdrawal, especially given the role of neurosteroid based treatments in managing several conditions with inflammatory involvement (Morrow et al. 2020). Neurosteroids have been shown to mediate alcohol related behaviors and buffer the impact of neuroinflammation (Fujii et al. 2021). We see an effect in acute withdrawal; while behavior is not altered by the 5α-reductase knockdown in the BLA in males or females, we do see a trend towards a negative correlation between allopregnanolone after CIE exposure and open field anxiety like behavior in 5α-reductase knockdown females but not males (Figure 5A-B). Higher levels of allopregnanolone may be associated with increased avoidance behavior in the knockdown animals and not the control virus animals (lower ratio of time in the open zone over the periphery). This is inconsistent with the literature given allopregnanolone typically exerts anxiolytic effects, but high levels of neuroinflammation could be resulting in altered or blunted effects of allopregnanolone. The increase in peripheral allopregnanolone levels may also represent a compensatory change associated with impaired neurosteroidogenesis in the brain. Future work is required to examine the relationship between inflammation, neurosteroids, and alcohol on withdrawal behaviors. Thus, the impact of allopregnanolone on outcome measures following chronic alcohol exposure are diverse and include anxiolytic effects as well as effects on neuroinflammation. Future work is required to fully understand the relationship between inflammation, neurosteroids, and alcohol on withdrawal behaviors.

One unanswered question relevant to these studies is how 5α-reductase expression is regulated. These studies demonstrate that 5α-reductase expression is decreased in the BLA following CIE. Similarly, our lab previously demonstrated that 5α-reductase expression is downregulated following chronic stress exposure (Walton et al. 2023). However, the mechanisms through which 5α-reductase becomes downregulated in response to stress or alcohol is unclear. This data suggests that 5α-reductase expression may be positively regulated by either steroid hormone precursors or the neurosteroids themselves. Additional studies are required to understand the regulatory mechanisms controlling endogenous neurosteroidogenesis which could have untapped therapeutic potential.

Our findings demonstrate sex differences in the interaction between neurosteroids and alcohol, suggesting a sexually dimorphic mechanism in how allopregnanolone is mediating ethanol consumption. Allopregnanolone is the focus of the current study, which is a metabolite of progesterone. However, 5α-reductase also plays a role in the metabolism of testosterone. Studies have linked testosterone and opioid misuse with expression levels of srd5α2 in human patients (Janke et al. 2021) as well as alcohol misuse in adolescents. Increased testosterone was associated with higher alcohol intake in boys acting via disrupted amygdala-orbitofrontal cortex connectivity (Peters, 2015). In addition, knockdown of 5α-reductase may have unintended impacts on the synthesis of other neurosteroids by preventing conversion to 5α-dihydroprogesterone and then allopregnanolone. The neurosteroidogenic enzymatic pathway is complex and involves the synthesis of neurosteroids which impact both NMDA receptors and GABA receptors. Thus, we cannot rule out impacts on other neurosteroids which may impact withdrawal behaviors and alcohol intake.

The current study is limited in that it could benefit from an investigation of how the estrous cycle and housing conditions impact ethanol consumption. While the female mice used in this study were acyclic, multiple studies have reported the impact of estrous cycle on ethanol associated behaviors and reward seeking (Brot et al. 1995, Dazzi et al 2007, Giacometti et al 2022). Given the synthesis of allopregnanolone from progesterone (Diviccaro et al. 2022) and clinical work showing allopregnanolone levels are sensitive to ovarian cycle phases (Tores and Ortega 2003) it is likely that endogenous levels of allopregnanolone and 5α-reductase may affect how animals respond to ethanol vapor, 5α-reductase knockdown, and chronic treatment with SGE-516 chow. Animals were also singly housed for drinking experiments which is a known stressor and could impact neurosteroid synthesis and their behavior in avoidance tasks. Ethanol consumption has been shown by Peterson et al. 2024 to be influenced by the housing environment; single housing amplified ethanol consumption in both male and female mice, which was reversed upon return to group housing. Future studies will consider this role on environment on the impact of allopregnanolone, stress and ethanol associated behaviors. This is particularly important given our findings that 5α-reductase expression is also impacted by chronic stress exposure (Walton et al. 2023) and exogenous allopregnanolone influences behavioral outcomes following chronic stress exposure (Antonoudiou et al. 2021).

Overall, we demonstrate that endogenous neurosteroidogenesis in the basolateral amygdala is impaired by chronic ethanol exposure. Importantly, these studies demonstrate that treatment with exogenous neurosteroid-based treatments may have unanticipated effects on alcohol consumption during the acute withdrawal period. Thus, this study is important in understanding how allopregnanolone synthesis in the BLA contributes to hyperkatifeia in withdrawal, especially since allopregnanolone is a promising treatment for patients in recovery for alcohol misuse disorder. It is also important to understand the sex differences in how allopregnanolone is interacting with the brain in the context of alcohol withdrawal.

## Acknowledgements

The authors would like to recognize the support and guidance for Dr. Klaus Miczek on this project. The authors thank Dr. Phil Haydon for the use of the La Jolla ethanol vapor chambers. This work was supported by a grant from NIAAA: R01AA026256 (J.M.), NIGMS: R25 HL007785 (J.M.), and NIMH: R01 MH128235 (J.M.).

## 5. Supplemental Data Figures

**Supplemental Figure 1.**
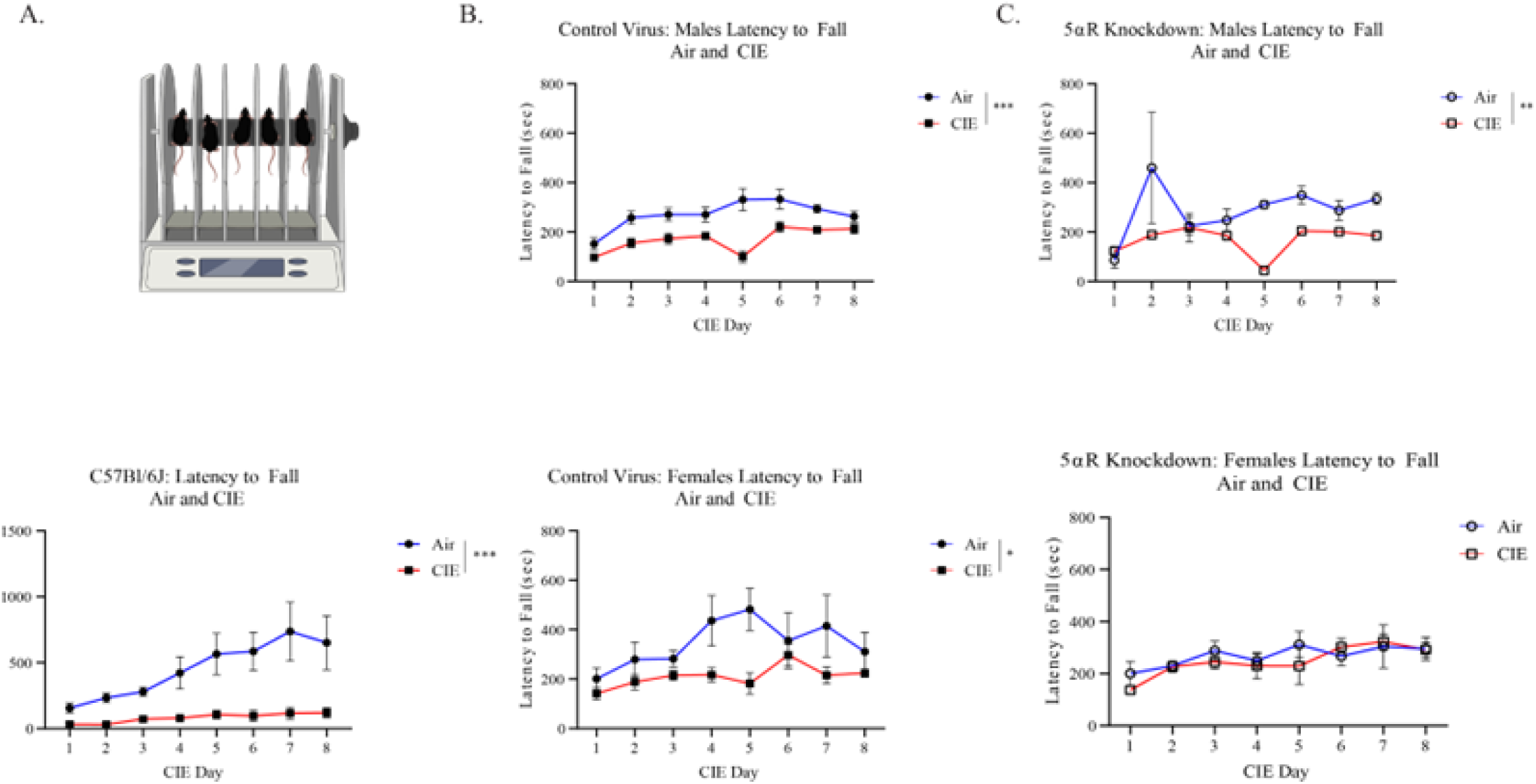
Latency to fall from the rotor rod is decreased in CIE mice with the exception of female 5_α_-reductase knockdown animals. A . Rotor rod latency to fall over the eight days of CIE in C57Bl/6J animals exposed to air ( n=10) and CIE (n=8-12, Mixed Effects Model, Tx History x Time, Effect of Tx History p=0.0001) B. Rotor rod latency to fall over the eight days of CIE in control virus CRISPR::Cas9 male Air and CIE mice (top, n=10 Air, 8 CIE, Mixed Effects Model, Tx History x Time, Effect of Tx History, p=0.0006) and female Air and CIE mice (bottom, n=4 Air, 8 CIE, 2-Way ANOVA Tx History x Time, Effect of Tx History, p=0.0460). C. Rotor rod latency to fall over the eight days of CIE in 5_α_-reductase knockdown male Air and CIE (top, n=7 Air, 12 CIE, Mixed Effects Model, Tx History x Time, Effect on Tx History, p=0.0018) and female Air and CIE mice (bottom, n=4 Air, 6 CIE, 2Way ANOVA Tx History x Time, Effect of Tx History, p=0.6174). \**p* <0.05, \*\**p*<0.01, \*\*\**p* <0.001 across all timepoints? Data are presented as the mean±SEM

**Supplemental Figure 2.**
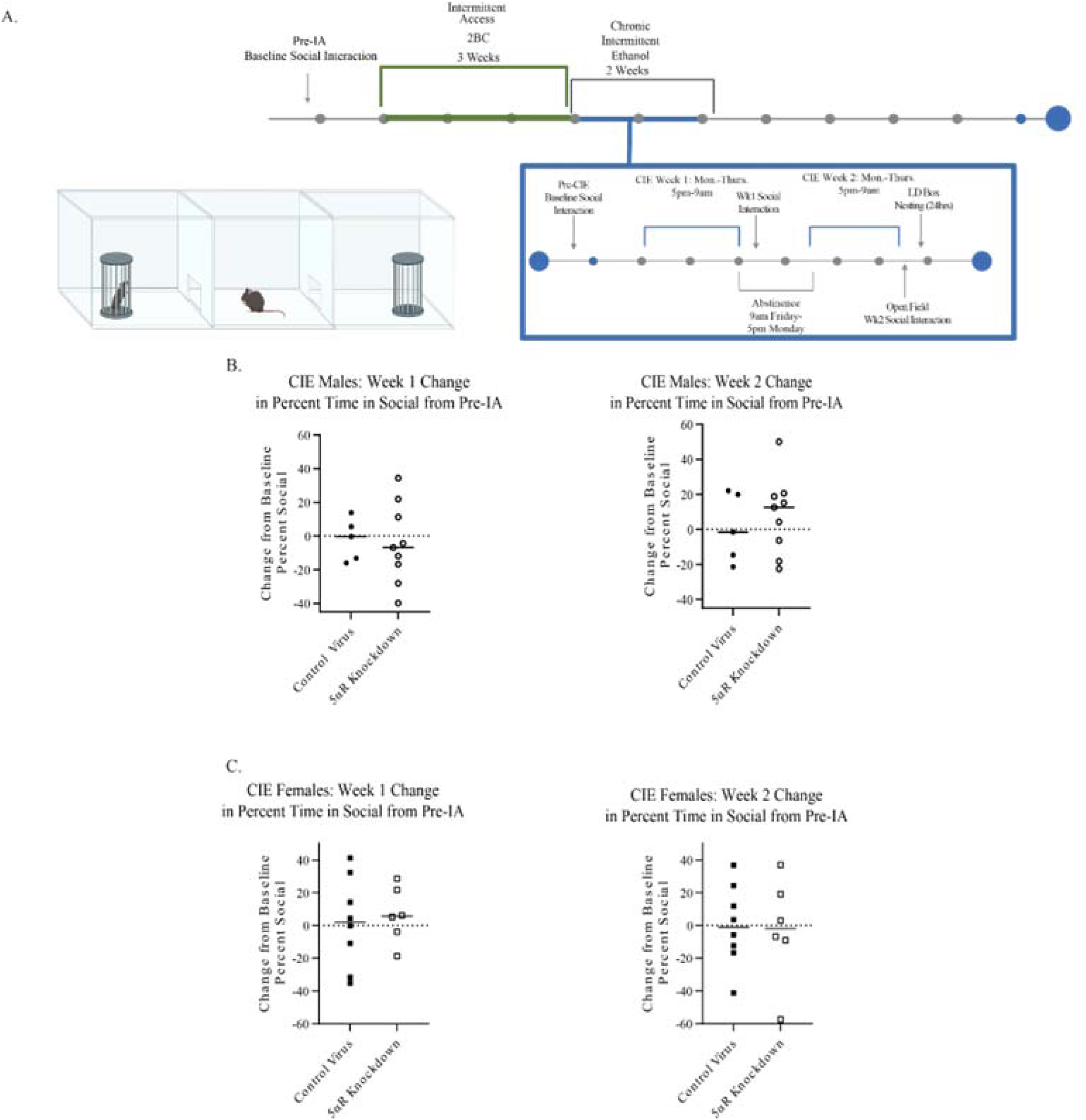
Social interaction is not impacted by a knockdown of 5_α_-reductase in males or females at week 1 and 2 of CIE exposure. A. Timeline of social interaction behavior testing. B. Change in percent time spent in social chamber from the initial pre-IA social interaction task in male CIE mice at CIE week 1 (left, n=5 control virus, 9 5αR knockdown, One Sample T tests, Theoretical Mean 0.0, control virus, p=0.7369, 5αR knockdown, p=0.5849) and at CIE week 2 (right, n=5 control virus, 9 5_α_R knockdown, One Sample T tests, Theoretical Mean 0.0, control virus, p=0.9272, 5_α_R knockdown, p=0.2998). B. Change in percent time spent in social chamber from the initial pre-IA social interaction task in female CIE mice at CIE week 1 (left, n=8 control virus, 6 5_α_R knockdown, One Sample T tests, Theoretical Mean 0.0, control virus, p=0.8608, 5_α_R knockdown, p=0.3930) and CIE week 2 (right, n=8 control virus, 6 5_α_R knockdown, One Sample t tests, Theoretical Mean 0.0, control virus p=0.9968, 5_α_R knockdown p=0.8625). Data are presented as the mean±SEM.

**Supplemental Figure 3.**
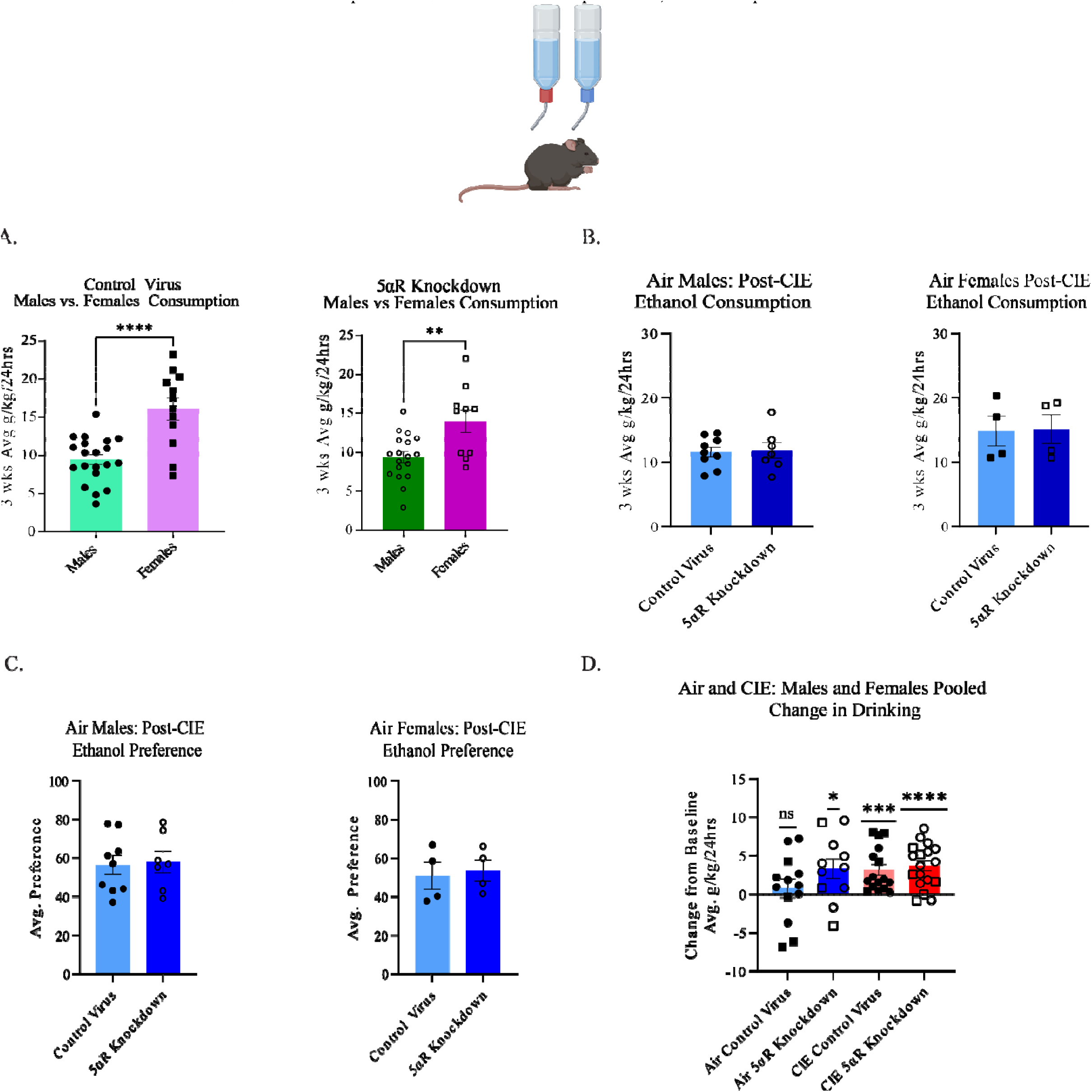
Females drink more than males at baseline in both groups and drinking is escalated from baseline after CIE in CIE animals and the Air 5_α_R knockdown animals. A . Baseline (pre-CIE) ethanol consumption in male and female control virus (left, n=20 males, 12 females, unpaired t-test, p<0.0001) and 5_α_R knockdown animals (right, n= 18 males, 10 females, Unpaired T test, p=0.0031). B. Ethanol consumption post-CIE in air treated males (left, n=9 control virus, 7 5_α_R knockdown, Unpaired T test, p=0.8603) and females (right, n=4 control virus, 4 5_α_R knockdown, Unpaired T test, p=0.9320). C. Ethanol preference post-CIE in males (left, n=9 control, 7 5_α_R knockdown, Unpaired T test, p=0.8313) and females (right, n=4 control, 4 5_α_R knockdown, Unpaired T test, p=0.7706). D. Change in drinking from baseline in males (circles) and females (squares) in air control virus (n=13,One Sample T test, Theoretical Mean 0.0, p=0.5260) air 5_α_R knockdown (n=11, One Sample T test, Theoretical Mean 0.0,p=0.0275), CIE control virus (n=16, One Sample T test, Theoretical Mean 0.0, p=0.0005) and CIE 5_α_R knockdown animals (n=18, One Sample T test, Theoretical Mean 0.0, p<0.0001). ns= not significant, \**p* <0.05, \*\*\**p*<0.001, \*\*\*\**p* <0.0001? Data are presented as the mean±SEM.

**Supplemental Figure 4.**
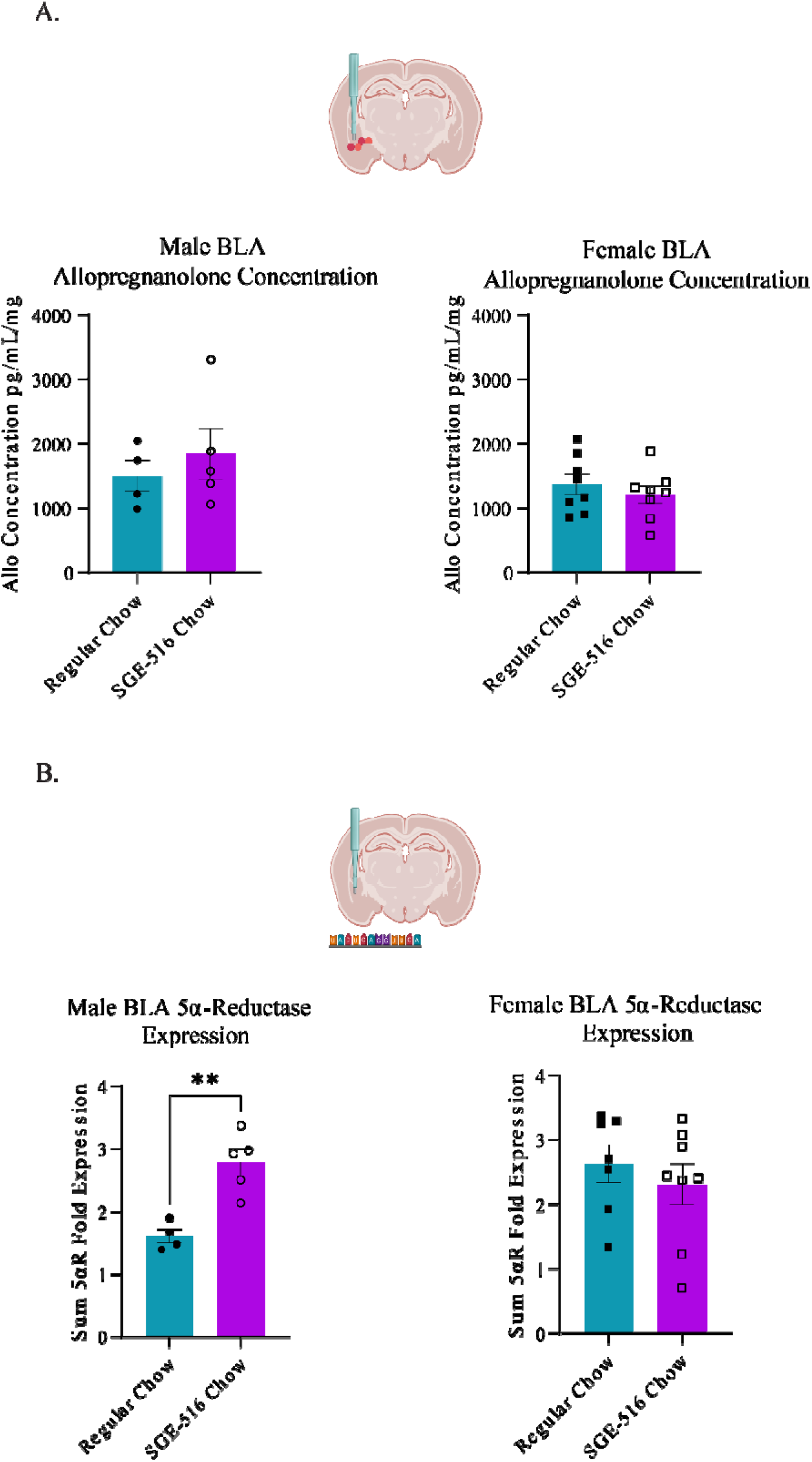
The capacity for neurosteroid synthesis is increased in male mice fed SGE-516 chow. A. Allopregnanolone in the BLA of male regular chow and SGE-516 chow (top, n= 4 regular chow, 5 SGE-516 chow, Unpaired T test, p=0.5086) and female regular chow and SGE-516 chow (n=8 regular chow, 8 SGE-516 chow, Unpaired T test, p=0.4608). B. 5α-reductase transcript levels in the BLA of male (top, n=4 regular chow, 5 SGE-516 chow, Unpaired T test, p=0.0028) and female mice (bottom, n=7 regular chow, 8 SGE-516, Unpaired T test, p=0.5503). Data are presented as the mean±SEM.

## Notes

### Competing Interest Statement

J.M. serves on the Scientific Advisory Board for Ovid Therapeutics for work unrelated to this study. This work was done in collaboration with SAGE Therapeutics who provided the SGE-516 compound, but did not fund this study. M.L. and S.G. were previously employed and J.M. previously served on the Scientific Advisory Board for SAGE Therapeutics. All other authors declare no conflict of interest. 

